# Efficient Prohibitin 2 exposure during mitophagy depends on Voltage-dependent anion-selective channel protein 1

**DOI:** 10.1101/2023.10.10.561633

**Authors:** Moumita Roy, Sumangal Nandy, Elena Marchesan, Chayan Banerjee, Rupsha Mondal, Federico Caicci, Elena Ziviani, Joy Chakraborty

## Abstract

Autophagic elimination of depolarized mitochondria (mitophagy) depends on Ubiquitin proteasome complex to expose the inner mitochondrial membrane-resident protein-Prohibitin 2 (PHB2). This uncovering facilitates its interaction with autophagosomal membrane-associated protein LC3. It remains unclear whether PHB2 is uncovered randomly through mitochondrial rupture sites. Prior knowledge and initial screening indicated that Voltage-dependent anion-selective channel protein 1 (VDAC1) might play a role in this process. Through *in vitro* biochemical assays and imaging, we have found that VDAC1-PHB2 interaction increases during mitochondrial depolarization. Subsequently, this interaction enhances the efficiency of PHB2 exposure and mitophagy. To investigate the relevance *in vivo*, we utilized a Porin (equivalent to VDAC1) knockout *Drosophila* line. Our findings demonstrate that during rotenone-induced mitochondrial stress, Porin is essential for PHB2 exposure, PHB2-LC3 interaction, and mitophagy. This study highlights that VDAC1 predominantly synchronizes efficient PHB2 exposure through mitochondrial rupture sites during mitophagy. These findings may provide insights to understand progressive neurodegeneration.

## Introduction

Autophagic degradation of mitochondria (mitophagy) is one of the ways by which defective mitochondria are eliminated ^1–3^. This is an essential process that upholds the quality of the mitochondrial population within the cellular environment ^4–6^. The presence of various routes to support mitophagy underscores the significance of this process in ensuring the survival of cells. ^7^. In this context, one of the extensively investigated pathways involves PTEN-induced kinase 1 (PINK1, a kinase) and Parkin (an E3 ubiquitin ligase)-mediated mitophagy ^1, 2^. Mutations in these proteins are closely associated with neurodegenerative conditions, notably Parkinson’s disease (PD)^8, 9^. It is unequivocally acknowledged that following depolarization, PINK1 recruits Parkin to the outer mitochondrial membrane (OMM), where Parkin subsequently ubiquitinates various target proteins. PINK1 can phosphorylate multiple outer mitochondrial membrane (OMM) proteins, with some of them serving as docking sites for Parkin, yet only a limited subset of OMM proteins have been identified as fulfilling this particular function. ^10, 11^. From an alternate standpoint, it is proposed that PINK1 might phosphorylate a pre-existing ubiquitin chain on OMM, subsequently initiating the recruitment of Parkin. As a result, the substrate undergoes further ubiquitination, thereby amplifying the process. Mitochondrial E3 ligases (like MITOL) are essential for such initial seeding effects, but in the absence of Parkin, they alone are insufficient to induce mitophagy. ^12, 13^. Up to this stage, the process can be partially reversed, as the ubiquitination mediated by Parkin is known to be counteracted by various deubiquitinase enzymes. ^14^. If ubiquitinated proteins on OMM are not subjected to deubiquitination, they can either be degraded by the Ubiquitin-proteasome complex or associate with P62. ^15, 16^. P62, serving as an adapter protein, enables the linkage between ubiquitinated proteins and Microtubule-associated proteins 1A/1B light chain 3B (LC3). Earlier research has indicated that the Ubiquitin-proteasome complex is equally vital in depolarization-induced mitophagy, although the precise rationale remained unclear ^17, 18^. Recently, it has come to light that mitophagy requires proteasome activity-dependent rupture of OMM and exposure of inner mitochondrial membrane (IMM) protein Prohibitin 2 (PHB2) ^19^. PHB2 can function as an LC3 receptor. Unlike Cardiolipin, which translocate to OMM during depolarization ^20^, OMM rupture site formation is important for such PHB2 binding with LC3 ^19^.

Prohibitins (PHB1 and 2) are membrane proteins that are highly conserved and exist as multimeric ring complexes within mitochondria. They are involved in several crucial processes, such as mitochondrial shaping, biogenesis, and maintenance of mitochondrial DNA ^21–23^. Notably, PHB2 has the intriguing capacity to control the release of Cytochrome c by safeguarding OPA1 long forms from mitochondrial proteases. Although cells lacking PHB2 can survive, they are highly sensitized to apoptotic stimuli ^23^. Recently it was revealed that the PHB complex’s scaffolding effect on PARL-PGAM5 can also regulate PINK1 processing ^24^. The significance of PHB2 in mitophagy is further emphasized by another study, which showed that Parkin can ubiquitinate PHB2 when OMM is disrupted. ^25^.

While several studies have investigated the PINK1-Parkin pathway up to the stage of OMM rupture and subsequent PHB2 exposure, none elucidated how PHB2 remains in close proximity to the sites of rupture. Here, we investigated two possible explanations: i) PHB2 exposure after mitochondrial depolarization is a random event or ii) PHB2 interacts with an OMM Parkin target, thus degradation of such a protein may increase the efficiency of PHB2 exposure. If this interacting protein on OMM could form oligomers (or form a multimer with other OMM Parkin targets), it may further enhance the efficiency.

We utilized an unbiased approach to identify a potential partner of PHB2 that could be targeted for ubiquitination by Parkin. Our study reveals that during stress-induced mitophagy, Voltage-dependent anion-selective channel protein 1 (VDAC1) can facilitate the exposure of PHB2 to LC3. This study enhances our understanding of the underlying mechanisms in mitophagy, and the implications are broad. For example our findings have the capacity to explain issues related to mitochondrial quality control in progressive neurodegenerative disorders.

## Results and discussion

### Association of VDAC1 with PHB2 increases following mitochondrial depolarization

The primary aim of our study was to identify possible candidates capable of interacting with both PHB2 and Parkin. We reasoned that proteasome mediated degradation of such a protein on the depolarized mitochondrial surface could potentially result in the exposure of PHB2. Figure S1A depicts different scenarios illustrating variations in PHB2 exposure following mitochondrial depolarization and the degree of interaction with LC3. The scenarios depicted are as follows: (1) PHB2 is randomly exposed after the degradation of the Parkin substrate, (2) the Parkin target can oligomerize, thereby augmenting PHB2 exposure, (3) the Parkin target can oligomerize, but PHB2 is not abundantly located near the rupture site, (4) the substrate oligomerizes and engages with PHB2, (5) the Parkin substrate does not undergo oligomerization but interacts with PHB2. Our hypothesis aligns with the notion that scenarios 2, 4, and 5 could potentially result in significantly greater PHB2 exposure. We utilized BioGrid open database to identify a total of 233 proteins that could interact with both Parkin and PHB2 (Figure S1B). Out of these, 88 are mitochondrial proteins and 18 of them localize on OMM (Mitocarta 2.0 database). Importantly, we conducted GEO analysis (from GEO data set GSE20333 and GSE20186) on these common interactors, and found that expression of VDAC1 was significantly lower in the substantia nigra (SN) region of PD brain (Figure S1C and D). This is noteworthy since compromised mitophagy is strongly linked with progressive neurodegeneration in PD. Next, mitochondria were isolated from rat brain and divided in two equal portions. In one portion mitochondria were depolarized with with CCCP (10 µM, 20 min). The other portion received equal amount of vehicle. We used protein concentrators with 50 or 100 kDa cut-off polyethersulfone membranes to segregate the mitochondrial proteins. VDAC1 has a molecular weight of ∼34 kDa. Hence, if VDAC1 forms complexes with proteins of a molecular weight ranging from 15-20 kDa or higher, these membranes will obstruct their passage. We observed a significantly higher retention of VDAC1 and VDAC2 by the 50 and 100 kDa cut-off membranes when mitochondria were depolarized, suggesting possible oligomer or complex formation (Figure S1E). We selected VDAC1 for further investigation due to its high abundance on OMM and its high affinity towards Parkin ^14^. CCCP treatment (10 µM) did not alter total level of VDAC1 in isolated rat brain mitochondria (Figure S1E).

Next, we wanted to reconfirm that VDAC1 can interact with PHB2. As VDAC1 and PHB2 share similar molecular weight (∼34kDa), to avoid false positive results we utilized WT or VDAC1 KO HEK cells. We transfected VDAC1-3xFLAG into the VDAC1 KO cells and performed immunoprecipitation for FLAG. After immunoblotting, when compared to the WT untransfected cells, a shift of VDAC1 signal was noted in VDAC1-3xFLAG samples (Figure S1F). This is due to the additional ∼3 kDa 3XFLAG attachment with VDAC1. The signal for VDAC1 also overlapped with that of FLAG. In the FLAG immunoprecipitated samples we detected PHB2 signal at similar positions when compared to the WT. This reconfirmed that VDAC1 can interact with PHB2. Protein samples from VDAC1 KO cells, which were immunoprecipitated for FLAG, did not show VDAC1/FLAG/PHB2 signal (Figure S1F).

Previously it was noted that VDAC1 exhibits weak associations with one another, making it challenging to detect oligomeric forms (or complexes with PHB2, in this instance) after SDS-PAGE unless chemically crosslinked ^26, 27^. At first, we treated isolated rat brain mitochondria with CCCP (10µM, 20 min) and prepared EGS crosslinked protein samples for SDS-PAGE. Immunoblot analysis showed prominent VDAC1 signals at ∼ 34, ∼72, ∼130 and ≥180 kDa (Figure 1A). Following CCCP treatment, we consistently observed an increase in VDAC1 signal at ∼ 130 kDa and ≥180 kDa (figure 1A). Strikingly, the signal patterns of PHB2 were similar to VDAC1 at ∼72, ∼130 and ≥ 180 kDa (Figure 1A). To investigate this phenomenon under endogenous conditions we utilized SH-SY5Y cell line. We found that Mitochondrial mass or VDAC1 levels did not reduce within 2h of CCCP (10 µM) treatment (Figure S2A and B). We did not find any notable OMM rupture within this time period of CCCP treatment. Interestingly increased contact points between OMM and IMM were visible (Figure S2C). We observed an increase in high molecular weight VDAC1 and PHB2 signals (at∼72, ∼ 130 and ≥180 kDa) after 2h CCCP treatment in SH-SY5Y cells (Figure 1B). To confirm whether the heightened co-localization signals of VDAC1 and PHB2 result from their interaction, we subjected isolated rat brain mitochondria to CCCP treatment (for 20 minutes) and subsequently conducted immunoprecipitation for either VDAC1 or PHB2. Likewise, SH-SY5Y cells were exposed to CCCP for 2 h, mitochondria were isolated and the experiment was repeated. In both scenarios, we observed an increased interaction between VDAC1 and PHB2 upon CCCP treatment (Figure 1C). For further confirmation, we designed a split GFP based VDAC1-PHB2 interaction sensor, where GFP moiety 1-10 was attached to VDAC1 C-terminal and GFP β strand 11 was attached to N-terminal end of PHB2. These two parts of GFP remained non-fluorescent (Figure S3). However, GFP can self-assemble and demonstrate prominent fluorescence when VDAC1 and PHB2 come in close proximity. VDAC1-GFP (1-10) and PHB2-GFP (11) were expressed in VDAC1 KO HEK cells. Control HEK cells (without CCCP) demonstrated puncta like GFP signals inside mitochondria (Figure 1D). 20 min CCCP treatment led to heighted GFP signal, signifying increased self-assembly of GFP and thus higher interaction between VDAC1 and PHB2 (Figure 1D).

**Figure 1.**
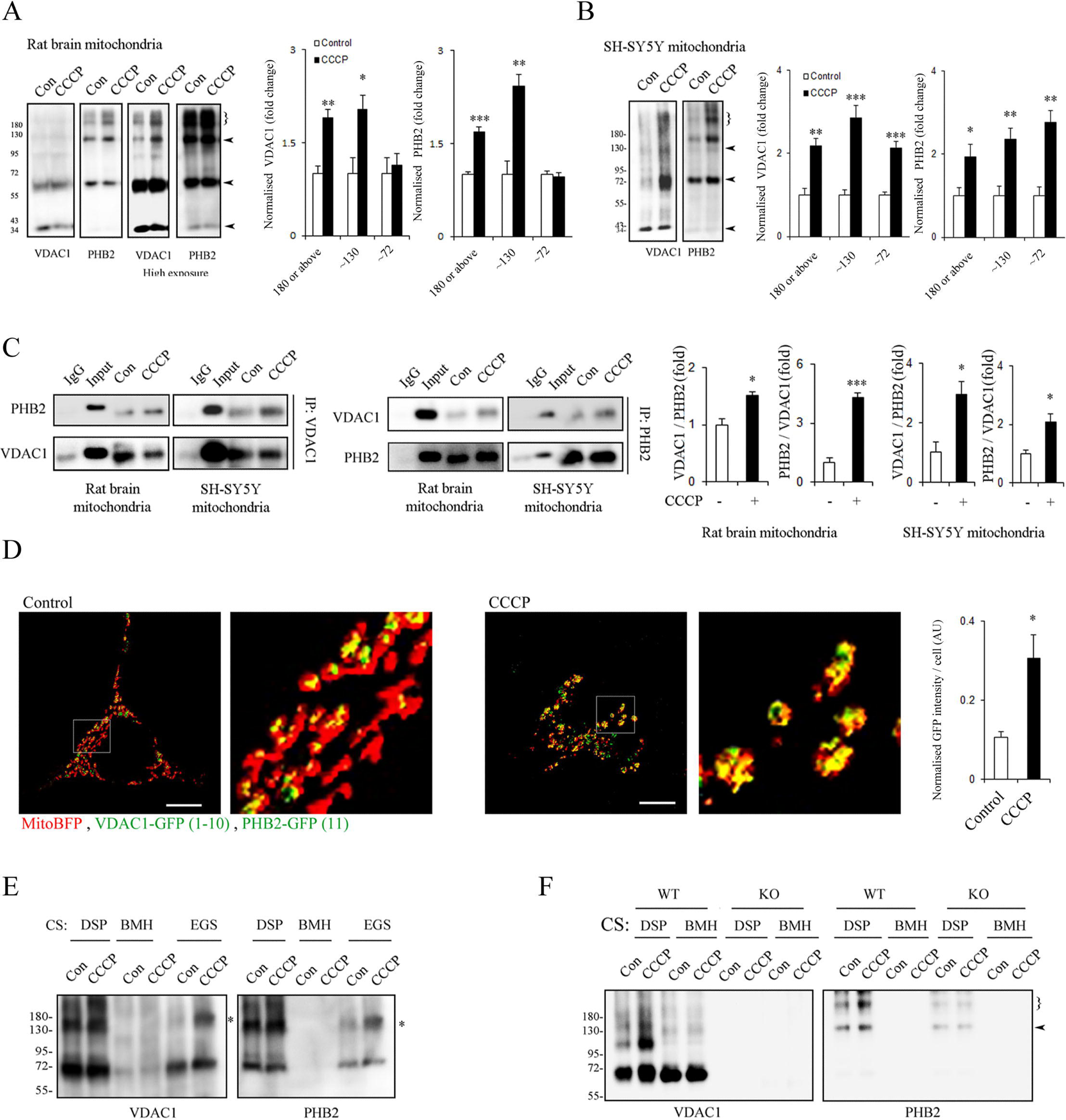
VDAC1-PHB2 complex formation during CCCP induced depolarization. **A.** Isolated rat brain mitochondria are treated with CCCP (10 µM) for 20 minutes, crosslinked (CS) with EGS and subjected to immunobloting. High and short exposure images of the same blot are provided. ∼70, ∼130 and ≥180 kDa band intensity were normalized by ∼34 kDa band intensity and represented as bar graphs. **B.** SH-SY5Y cells are treated with CCCP (10 µM) for 2h and isolated EGS CS mitochondria are processed for immunobloting. ∼70, ∼130 and ≥180 kDa band intensity were normalized by ∼34 kDa band intensity and represented as bar graphs. **C.** Isolated rat brain mitochondria are treated with CCCP (20 min), or isolated mitochondria from 2h CCCP treated SH-SY5Y cells are immunoprecipitated for either VDAC1 or PHB2. After SDS-PAGE, immunoblots are probed with the indicated antibodies. **D.** VDAC1 KO HEK cells transfected with VDAC1-GFP (1-10), PHB2-(GFP11) and mitoBFP are treated with vehicle or CCCP (10 µM, 20 min). 3D rendered z stack images are represented to demonstrate increased GFP signal after CCCP treatment. The experiment was repeated for 3 times and at least 45 cells were considered. Scale bar - 10 µm. GFP intensity was normalized by mitoblue intensity and mean values are represented as the bar graph. **E.** Rat brain mitochondria are treated as mentioned in A and crosslinked with DSP/BMH/EGS. Immunoblots represent VDAC1 and PHB2 localization after SDS-PAGE. (*) indicate the missing signal of VDAC1 in the BMH crosslinked samples. **F.** Mitochondria isolated from control or VDAC1 knockout (KO) HEK293T cells are treated as mentioned in C and crosslinked as mentioned in figure. Immunoblots are probed with VDAC1 and PHB2 antibodies. Bar graphs represent mean fold change (± SEM). Student’s t test. *P≤ 0.05, **P ≤ 0.01, ***P ≤ 0.001 when compared to control group.

### The VDAC1-PHB2 complex constitutes the central element within the larger molecular assemblies

Next we wanted to determine whether this altered signal of VDAC1 at ∼130 and ≥180 kDa is dependent on the presence of PHB2 or not. Knockdown of PHB2 can alter mitochondrial morphology, polarization and membrane architecture ^22, 23, 28^. Given that mitochondria are already compromised, the impact of CCCP will not be decipherable in a PHB2 null background. So, we utilized the fact that rat PHB2 does not have cysteine residue, and thus should remain insensitive to BMH crosslinking. Although CCCP treated samples exhibited increased VDAC1 or PHB2 signal at ∼130 and ≥180 kDa after DSP or EGS crosslinking, no such effect was found after BMH crosslinked samples (Figure 1C). Rat VDAC1 contains two cysteines, thus VDAC1 signal at ∼72 kDa in BMH crosslinked samples represent VDAC1 dimer lacking any PHB2. Interestingly, VDAC1 signal at ∼130 kDa was not visible in BMH crosslinked samples, indicating the necessity of PHB2 to form the complex at ∼130 kDa (Figure 1C).

To determine whether increase in PHB2 signal at ∼130 and ≥180 kDa after CCCP treatment is VDAC1 dependent or not, we used VDAC1 KO HEK293T cells (Figure S4). The total amount of PHB2 does not change in absence of VDAC1 (Figure S4A-C). We isolated mitochondria from VDAC1 KO HEK cell lines and treated with CCCP (10 µM, 20 min). The inter-reliant alteration of PHB2 and VDAC1 signals (at ∼130 and ≥180 kDa) due to depolarization was visible in mitochondrial samples which were isolated from VDAC1 WT HEK293T cells (Figure 1D). Although DSP crosslinked protein samples from WT cells showed increased VDAC1 signal at ∼130 and ≥180 kDa after CCCP treatment, BMH crosslinking demonstrated no such effects. On the other hand, increased PHB2 signals were detected (at ∼130 and ≥180 kDa) when protein samples from WT cells were crosslinked with DSP. As anticipated, PHB2 signal was not detectable when BMH was used for crosslinking.

Interestingly, PHB2 signals did appear after DSP crosslinking at ∼130 and ≥180 kDa even when VDAC1 was absent. However, the CCCP induced increased signal at ∼130 and ≥180 kDa were not observed. The PHB2 signals observed at the same locations in the absence of VDAC1 might be a result of PHB2 binding to other proteins, and these complexes remained unaffected by mitochondrial depolarization.

In order to ascertain if VDAC1 oligomerization is necessary for the formation of the ∼130 kDa complex, we pre-treated rat brain mitochondria with VDAC1 oligomerization inhibitor-DIDS ^26, 27, 29^. DIDS effectively inhibited the complex formation of VDAC1 at ∼130 kDa in response to CCCP treatment (Figure S5A). If PHB2-VDAC1 (with a combined molecular weight of ∼ 68 kDa) serves as the fundamental component of ∼130 kDa or larger complexes, the breakdown of these complexes should result in signals corresponding to their individual molecular weights (∼34 kDa). Complete disintegration of DSP crosslinked samples by β mercaptoethanol exhibited VDAC1 and PHB2 signals only at ∼34 kDa (Figure S5B) in immunoblots after 2D SDS-PAGE. After first dimension SDS-PAGE, partial reduction of EGS crosslinked samples demonstrated clear signals of VDAC1 and PHB2 at ∼ 34 and ∼72 kDa (Figure S5C). Smear like signals above 72 kDa represent intact or partially disintegrated complexes. Further reduction of ∼72 kDa complexes (indicated by rectangle, Figure S5B) yielded signals of VDAC1 and PHB2 at ∼34 kDa (Figure S5C).

### VDAC1 is required for PHB2 exposure and efficient mitophagy after CCCP treatment

Subsequently, we investigated whether VDAC1 KD, KO or inhibiting oligomerization could impede PHB2 exposure and mitophagy. Absence of VDAC1 does not interfere with depolarization induced Parkin localization to mitochondria (Figure S6). We monitored mitophagy at four different stages as mentioned in Figure S7A. We employed protease (Trypsin) protection or digestion assay to assess the integrity of OMM. Figure S7B demonstrates three different scenarios where Trypsin has differential access to TOM20 (OMM), PHB2 (IMM), ATP5a (IMM-cristae) and HSP60 (matrix), when incubated for a specific time period and at a specific temperature. Trypsin digestion assay in SH-SY5Y cells revealed that ATP5a, is partially digested by trypsin after 4 hours of CCCP treatment. This might indicate the beginning of robust OMM breakdown (Figure S7C and S7B-second scenario)(Wei, et al. 2017). Co-treatment of proteasome inhibitor MG-132 protected against CCCP-induced PHB2 exposure at 4h (Figure S7D and E). It was demonstrated that VDAC1 oligomers can form pores big enough to release mtDNA (macro-pore) (Kim, et al. 2019a). These macro-pores might uncover PHB2 situated near VDAC1 oligomers, without requiring OMM rupture by proteasome (Figure S7B - third scenario). Treating isolated mitochondria with CCCP did not increase the sensitivity of PHB2 or other inner components like ATP5a and HSP60 towards Trypsin (Figure S7F). Thus, PHB2 exposure through VDAC1-macro pore formation was not perceived. After 4h CCCP treatment, PHB2 sensitivity towards Trypsin demonstrated that OMM is disintegrated in SH-SY5Y or HEK cells (Figure 1G and S8A). Interestingly, PHB2 is not efficiently exposed to Trypsin after CCCP treatment in VDAC1 KD or KO cells when compared to the control (Figure 2A). DIDS treatment also inhibited CCCP treatment induced PHB2 exposure in SH-SY5Y cells (Figure 2A).

**Figure 2.**
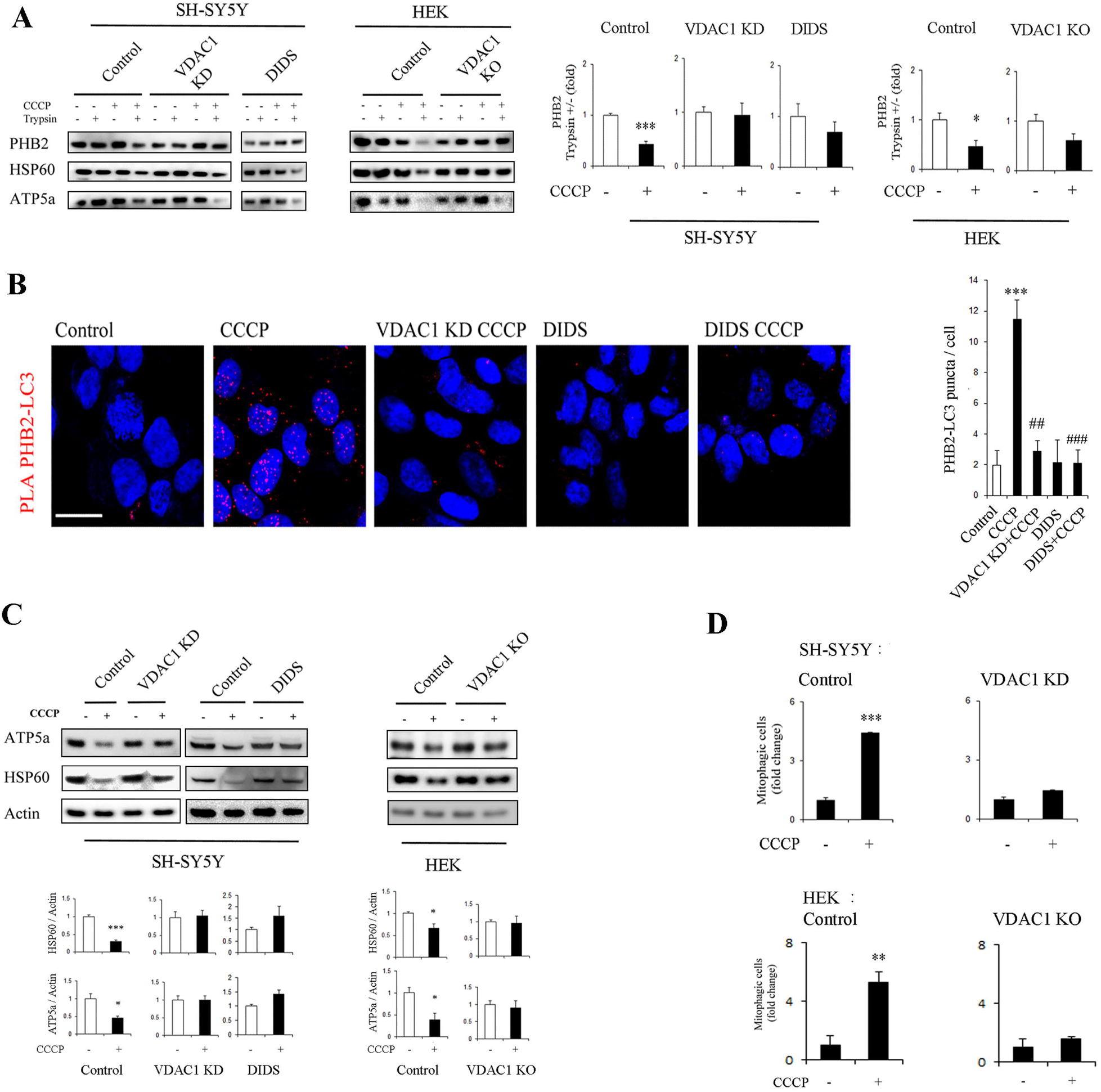
VDAC1 is required for depolarization induced PHB2 exposure and mitophagy. **A.** Immunoblots represent protease protection assay for PHB2, mitochondrial inner membrane (ATP5a) and matrix (HSP60) markers in the cell lines after the mentioned treatments. Cells were incubated with CCCP (10 µM) / DIDS (100 µM) for 4h. Bar graphs represent mean fold change (± SEM). *P ≤ 0.05, ***P ≤ 0.001 when compared to control group. N=3, Student’s t test. **B.** Representative images of Duo link proximity ligation assay for PHB2 and LC3 in SH-SY5Y cells after the mentioned treatments. Cells are treated with DIDS (100 µM) and / or CCCP (10 µM) for 4h. Scale bar 10 µm. Bar graphs represent mean number of PLA puncta (± SEM) / cell. ***P ≤ 0.001 when compared to control group. ^##^P≤ 0.01 / ^###^P≤ 0.001 when compared to the only CCCP treated group. The experiment is repeated three times. One way ANOVA followed by Tukey’s multiple comparisons test. **C.** SH-SY5Y or HEK293T cells are treated with CCCP for 8h. DIDS is co-treated with CCCP in the mentioned group. Immunoblots for the mentioned proteins are representative of at least three different experiments. Bar graphs represent mean fold change ± SEM. N=3, *P ≤ 0.05, ***P ≤ 0.001, Student’s t test. **D.** Quantification of mitophagic cells after tranfecting mito-keima is performed by FACS analysis. Bar graphs represent mean fold change (± SEM) in number of high mitophagic cells after 8h of CCCP (10 µM) treatment in the mentioned cell lines. The experiment is repeated 3 times. **P ≤ 0.01, ***P ≤ 0.001, Student’s t test.

Proximity ligation assay showed that the interaction between PHB2 and LC3 is dependent on the presence and oligomerization of VDAC1 (Figure 2B). We also determined the level of ATP5a and HSP60 (which reflect mitochondrial mass) in different experimental conditions. Clear decrease in ATP5a or HSP60 was noticed after CCCP treatment, in control SH-SY5Y or HEK cells. However, VDAC1 KD/KO or DIDS treatment inhibited this decrease in mitochondrial mass (Figure 2C). Furthermore, the number of cells with high level of mitophagy was quantified after transfecting the mentioned cell lines with mitokeima (Figure 2D). We found that VDAC1 was necessary for the increase in high mitophagic cells in response to CCCP treatment (Figure 2D).

Notably, we discovered that VDAC1 oligomerization is as crucial for mitophagy as the mere existence of VDAC1. However, it is not clear why a single molecular interaction between VDAC1 and PHB2 is not sufficient in this scenario. It is possible that an elevated quantity of VDAC1 in the complex, and its subsequent degradation by the proteasome complex, could lead to larger rupture sites on the OMM (macro-slit). This might potentially increase the effectiveness of PHB2-LIR domain exposure (as suggested in Figure S1A, 4th column).

### Porin-PHB2 complex formation increases during rotenone induced mitophagy, *in vivo*

To investigate the role of VDAC1 in mitophagy within an *in vivo* setting, we employed *Drosophila* as a model organism. *Drosophila* has been extensively used to delineate mitochondrial elimination pathways and as CCCP cannot be fed to the flies, many of these studies administered rotenone to induce mitophagy ^1, 30, 31^. Similar to previous studies (Kim, et al. 2019b), we also found that 1 mM rotenone exposure can induce mitophagy, as assessed in live wings or thoracic muscle of mitokeima expressing flies (Figure 3A). Results obtained from fixed thoracic muscles in ATG8a-mcherry expressing flies confirmed our findings, where rotenone treatment led to an increase in mcherry-ATG8a, LC3, and ATP5a overlapping signals (Figure 3B). TEM images also confirmed the existence of mitophagic structures in fly thorax following two days of rotenone treatment (Figure 3C). To determine whether rotenone induced mitochondrial stress can also lead to increased Porin (equivalent to mammalian VDAC)-PHB2 complex formation, we isolated mitochondria from control and rotenone-treated *Drosophila* and crosslinked with EGS for immunoblotting. As seen in our *in vitro* experiments, rotenone treatment increased Porin and PHB2 signals at ∼130 and ≥180 kDa (Figure 3D). Immunoprecipitation of Porin or PHB2 from isolated *Drosophila* mitochondria likewise revealed increased interaction between these proteins when flies were exposed to rotenone (Figure 3E and S8).

**Figure 3.**
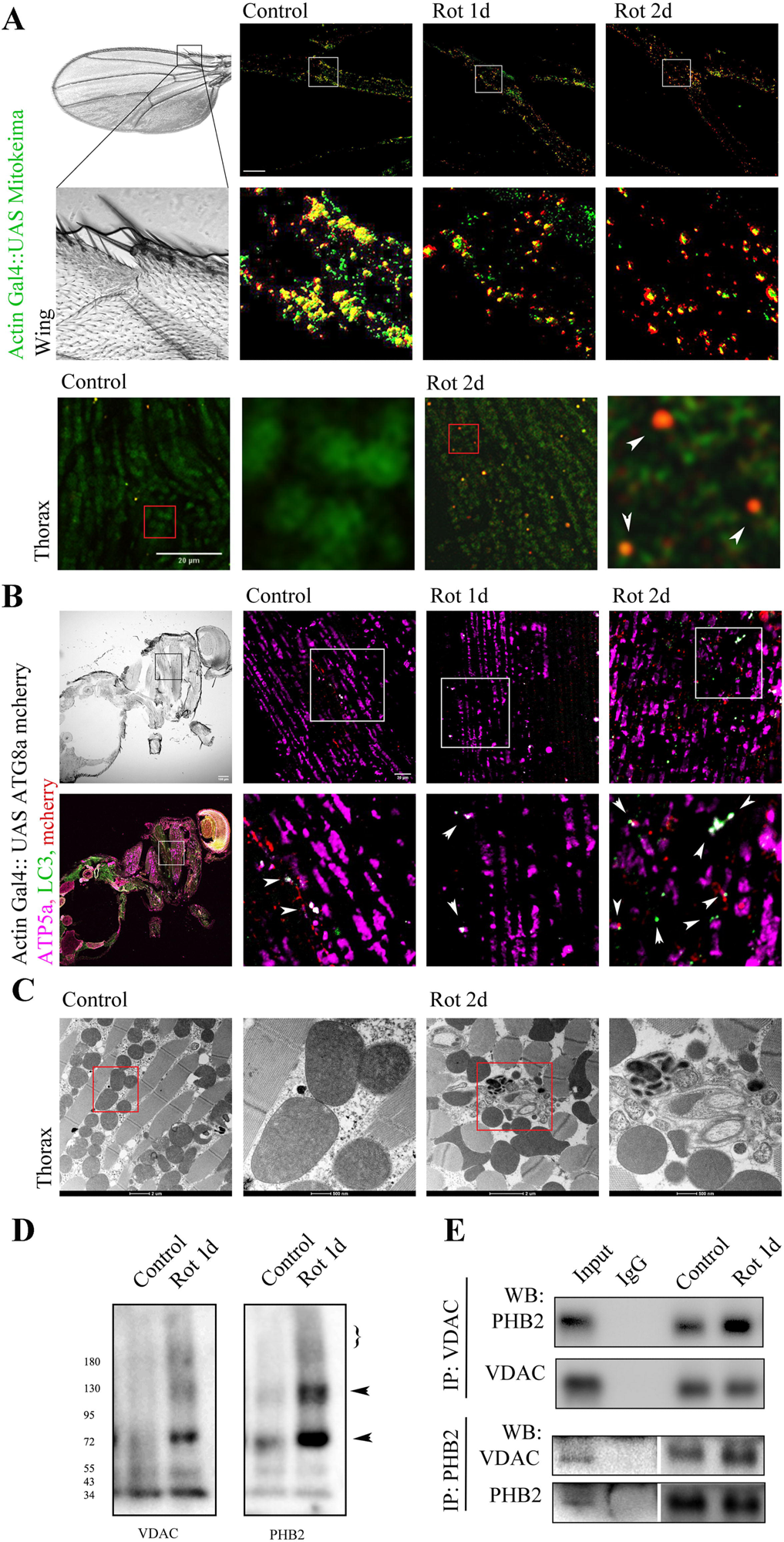
Rotenone treatment induces mitophagy and increases Porin-PHB2 complex formation in *Drosophila*. **A.** Images showing mitokeima signal from live fly wing (Actin Gal4: UAS mito Keima) after 1d or 2d rotenone (1 mM). For wing, images are taken from the indicated region in bright field image (scale bar 10 µm). Images for Mitokeima were also taken from live thoracic muscle after 2d of rotenone treatment. For Thoracic muscle mitochondria, we dissected the thorax and isolated muscles are imaged after submerging in phosphate buffer saline, pH 7.0 (scale bar 20 µm). Arrowheads indicated mitophagic structures. **B.** *Drosophila* expressing ATG8a-mcherry (Actin Gal4:UAS ATG8a-mcherry) are anaesthetized after 1d or 2d of rotenone treatment. Fixed fly thorax is immunostained for ATP5a, mcherry and LC3. Images are captured from the indicated portion of thorax as mentioned in bright field or fluorescent image of the whole body (scale 100 µm) and the representative merged images are provided. Arrowheads indicate colocalization of mcherry, ATP5a and LC3 signals, which appeared as bright white puncta (scale bar 20 µm). **C.** Fixed unstained thorax samples are processed for transmission electron microscope imaging and the selected regions from the representative images demonstrate presence of clear mitophagic structures after 2d of rotenone treatment (1 mM). Scale bar is as mentioned in the images. **D.** Isolated mitochondria from control and rotenone treated *Drosophila* are crosslinked with EGS and immunoblotted for VDAC1 and PHB2 as mentioned previously. Arrowheads indicate the coinciding signals. (}) indicate signals which appeared above 180 kDa molecular weight marker. Immunoblots are representative of three different experiments. **E.** *Drosophila* treated with rotenone (1d) are anaesthetized and mitochondrial protein samles are processed for immunoprecipitation. At least 20 flies are processed for individual experiment. Representative blots demonstrate signal for the mentioned proteins after co-immunoprecipitation (IP) for either VDAC or PHB2. Immunoblots are representative of three different experiments.

### Porin is required for rotenone induced PHB2 exposure and mitophagy in *Drosophila*

*Drosophila* has four different isoforms of VDAC, out of which Porin shows the highest similarity with mammalian VDAC1. Although Porin and Porin2 have similar functions, Porin is expressed ubiquitously whereas Porin 2 is restricted mostly to sperm cells ^32, 33^. To determine whether Porin is required for rotenone induced OMM rupture, PHB2 exposure and mitochondrial elimination, *in vivo*, we utilized WT, Porin KO (A2/A2) and Porin reintroduced/revertant (RV) *Drosophila* lines (Figure S9A)^34^. Thoracic mitochondria of A2/A2 flies appeared different from WT or RV lines (Figure S9B). We treated these flies with rotenone for 1day and mitochondria were isolated. When exposed to Trypsin, PHB2 displayed sensitivity in WT or RV fly lines (Figure 4A). In A2/A2 flies, PHB2 was not degraded by Trypsin (Figure 4A). Proximity ligation assay for LC3 and PHB2 showed increased proximity after rotenone treatment in the thoraces of WT and RV Drosophila. However, there were no significant changes detected in the thorax of A2/A2 flies (Figure 4B). Confocal microscopic images also revealed the presence of increased mitophagic structures in the thorax of WT and RV flies following two days of rotenone treatment (Figure 4C and S9C). TEM images confirmed the presence of mitophagic structures in these flies (Figure 8C). No clear alterations are observed in A2/A2 fly thorax (Figure 4C and S9C). Immunoblot analysis showed a decrease in ATP5a and HSP60 in WT or RV flies after a two-day rotenone treatment. There were no significant changes observed in A2/A2 flies (Figure 4D).

**Figure 4.**
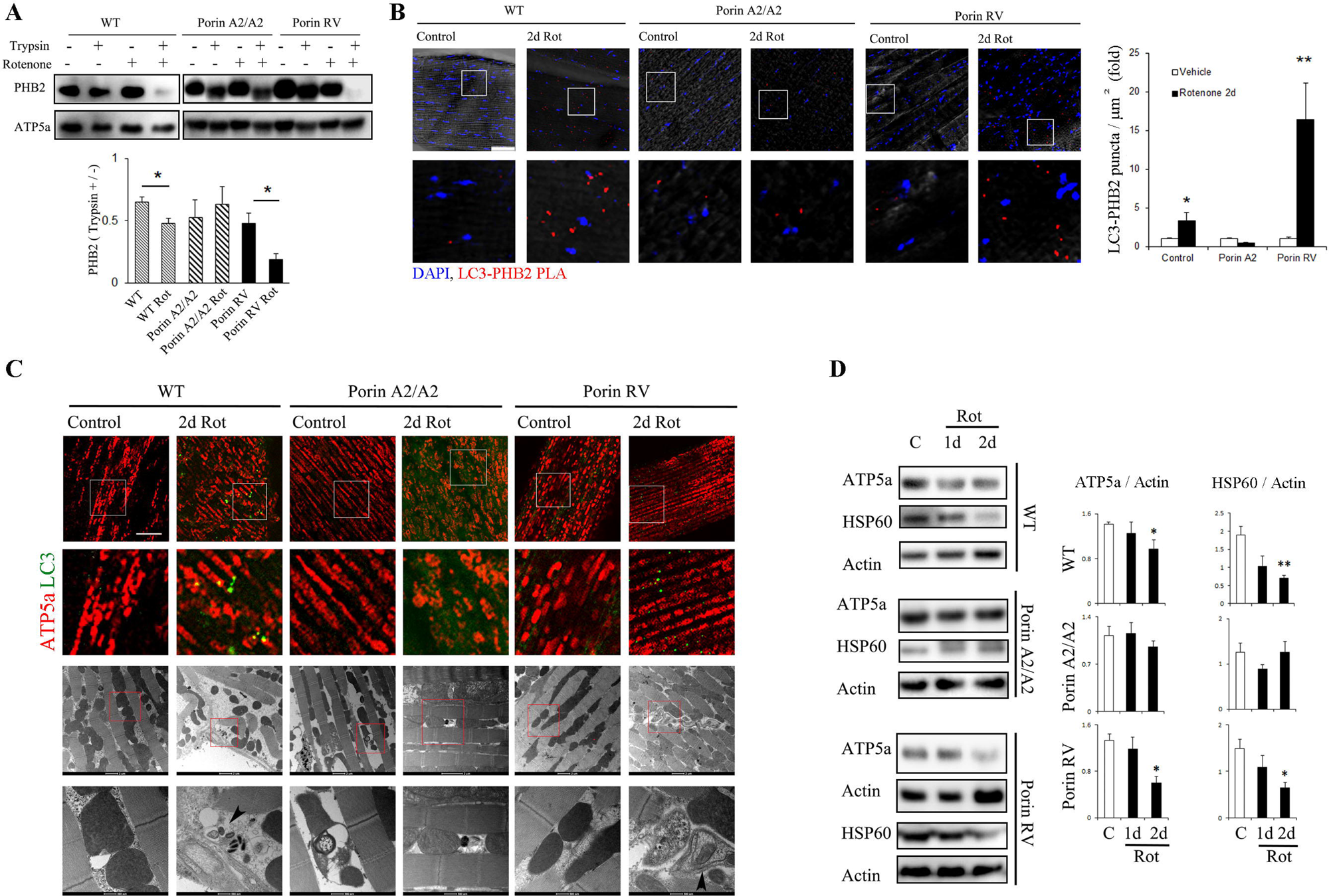
VDAC/Porin is necessary for rotenone induced mitophagy in *Drosophila*. **A.** Blots demonstrate protease protection assay of PHB2 in mitochondria isolated from WT, Porin KO (Porin A2/A2) and Porin revertant (Porin RV) *Drosophila* lines, after 1 day of rotenone treatment. Bar graphs represent mean ± SEM. *P ≤ 0.05. n=3-4, Student’s t test. **B.** Representative images of Duo link proximity ligation assay (PLA) for PHB2 and LC3 (red) in control or rotenone (2d) treated fly thorax of the mentioned lines are provided. Phase contrast and DAPI images are merged with the PLA signals. Bar graphs represent mean number of PLA puncta (± SEM). *P ≤ 0.05, **P ≤ 0.01. n=4-6. Student’s t test. **C.** Fixed thoracic samples of mentioned fly lines after 2d of rotenone treatment are processed for immunostaining or for transmission electron microscopy. Confocal microscope images represent colocalization of ATP5a (red) and LC3 (green). Scale bar 10 µm. Arrowheads in transmission electron microscopic images indicate mitophagic structures. Scale bar as mentioned in the image. At least 5 fly thoraces are imaged. **D.** Mitochondrial marker proteins (ATP5a and HSP60) from *Drosophila* lines are quantified by immunoblotting after 1d and 2d of rotenone treatment. Bar graphs represent mean intensity (± SEM). *P ≤ 0.05, **P ≤ 0.01. n=3-4. One way ANOVA followed by Dunnett’s multiple comparisons test.

Multiple research groups have reported that the activity of proteasome complex is required for breakdown of OMM at discrete sites during mitophagy. Discovery of LC3 receptor PHB2 exposure through these sites is relatively new and thus the efficiency of this unmasking process requires further characterizations. In this study, we report that the interaction between VDAC1 and PHB2 is crucial for effectively exposing PHB2 during depolarization or stress-induced mitophagy. The impact of this complex on mitophagy is also VDAC1 oligomerization dependent. Therefore, we propose that the degradation of oligomeric state of VDAC1 leads to the breakage of the OMM, creating a sufficient opening to expose PHB2. Our immunoprecipitation and protein crosslinking-based immunoblot assays suggest that VDAC1-PHB2 (∼64 kDa) may serve as the fundamental unit, while a ∼130 kDa component represents the simplest form of this complex which is relevant to PHB2 exposure and mitophagy. Importantly, our findings are applicable in an *in vivo* system where rotenone treatment with leads to a similar increased interaction between PHB2 and Porin. Additionally, the presence of Porin is necessary for unmasking of PHB2 and mitophagy in *Drosophila*, confirming the reproducibility of our in vitro findings. It is important to note that we employed CCCP and rotenone to induce stress in the system and investigate the influence of the VDAC1-PHB2 complex in enforced mitophagy. Whether this interaction remains significant in the context of normal physiological mitophagy during aging will necessitate further investigation.

An area that remained open for further study is whether the formation of this complex is specific to mitophagy or it serves as a broader response to various forms of mitochondrial damage. VDAC1 oligomerization has been frequently observed in the context of apoptosis induction ^27, 29, 35, 36^. The idea projects multimeric VDAC1 complex as a gateway to release mitochondrial inner compartment components, especially Cytochrome c. Conversely, VDAC1 is dispensable for the formation of mitochondrial permeability transition pore and VDAC1/3 knockout does not reduce apoptosis ^37^(Kim, et al. 2019a). This highlights that VDAC1 oligomerization and apoptosis might be stimulus and cell type specific (Kim, et al. 2019a). However, VDAC1 oligomeric states on apoptotic OMM are not well characterized. Little is known about other potential interacting partners of VDAC1 that may contribute to the formation of these complexes in diverse ways, particularly in response to various mitochondrial stressors.

A prior study suggests that VDAC1 plays a pivotal role at the intersection of mitophagy and apoptosis. The nature of VDAC1 ubiquitination is one of the determining factors, where mono-ubiquitination inhibits apoptosis and poly-ubiquitination leads to mitophagy (Ham, et al. 2020). This study also points out the significance of the PINK1-Parkin pathway in cell survival. During mitochondrial stress, there exists a continual competition between mitophagy and the possible release of Cytochrome c. Effective elimination of damaged mitochondria through mitophagy plays a crucial role in determining the outcome. A recent study has indeed shown that PINK-Parkin-induced proteasome-dependent rupture of the OMM can lead to the release of Cytochrome c when mitochondria are not adequately cleared through autophagic processes^38^. From our present findings, it appears that in situations where cellular autophagy mechanisms are active, the establishment of the VDAC1-PHB2 complex and the efficient exposure of PHB2 may have the potential to promote cell survival by facilitating mitophagy.

The question of whether the increased PHB2-VDAC1 interaction can offer protection against the impediments to mitophagy and the subsequent development of progressive neurodegenerative conditions (particularly in PD) warrants further examination. However, given its role in cell line and stimulus specific apoptosis, increased VDAC1 oligomerization may also lead to pathogenic conditions. Systemic lupus erythematosus (SLE) pathogenesis has recently been linked to elevated VDAC1 oligomerization ^26^. Although movement disorders are commonly associated with SLE, the occurrence of PD in such patients is relatively rare. Indeed, a sizable cohort study that assessed the likelihood of PD development in SLE patients discovered that these individuals have a reduced risk of PD occurrence ^39^. Further investigation is needed to determine whether the elevated VDAC1 oligomers associated with SLE include PHB2 as part of the complex, potentially promoting mitophagy for protection against PD development. It is worth noting that previous evidence has also shown reduced VDAC1 levels in the SN of PD brain ^40^(GEO database NCBI; accession number: GSE20333 and GSE20186).

Additional investigations and clarifications are needed to identify other proteins that interact with Parkin on the OMM. This will help determine if their association with PHB2 contributes to its exposure. Among the potential candidates, Mitofusin (1/2) and elements of the TOM complex stand out. TOM20, a component of the TOM complex, is known to form high molecular weight assemblies during pyroptosis ^41^. Nonetheless, the study provided evidence that mitochondrial depolarization induced by CCCP does not have the same impact. Mfn2 has not yet been established as an interactor with PHB2, particularly in the context of CCCP treatment. However, the elimination of Mfn2 does result in impaired mitophagy and retrograde neurodegeneration, possibly owing to its involvement in recruiting Parkin to the OMM ^42, 43^. These findings do not exclude the possibility that other roles of Mfn2, which are affected, may play a significant role in the development of Parkinson’s disease-like symptoms, such as the tethering between the endoplasmic reticulum and mitochondria ^44^.

It is important to note that while VDAC1 is a prominent substrate for Parkin, there is a scarcity of research dedicated to identifying factors that might influence VDAC1’s involvement in mitophagy ^14, 15, 17^. To summarize, our study demonstrated how VDAC1 can interfere with mitophagy by regulating PHB2 exposure towards LC3. While it is possible to bypass the need to degrade ubiquitinated proteins on OMM for mitophagy, exposure of PHB2 can increase the efficiency of the process. Thus, it can act as a critical factor in the development of neurodegenerative disorders. Further investigations are needed to delve into the connection between neurodegeneration and elements that impede the interaction between VDAC1 and PHB2.

## Materials ad Methods

### Reagents

All the reagents are purchased from Sigma-Alrich, unless mentioned otherwise.

### Animal ethics

Animal experimentation was done following the national guidelines (Care and Use of Animals in Scientific Research) formed by Committee for the Purpose of Control and Supervision of Experiments on Animals (CPCSEA), Animal Welfare Division, Ministry of Environment and Forests, Govt. of India. The protocol was evaluated and accepted by animal ethics committee of CSIR-Indian Institute of Chemical Biology, Kolkata, India.

### Antibodies and reagents

The following primary antibodies are used: VDAC1 (Millipore and Abclonal), VDAC, VDAC2, VDAC3, TOM20, ATP5a, Anti mCherry (Abcam), PHB2 (Sigma Aldrich), HSP60 (Biobharati, India), Actin (Santacruz Biotechnology), LC3 (Abcam and Abclonal). HRP and Alexa flour conjugated secondary antibodies are procured from Biobharati (India) and Invitrogen respectively.

All the other reagents are purchased from Sigma –Aldrich, otherwise sources are specified when mentioned.

### Plasmids

The following plasmids are used for the study: Parkin-YFP (Ziviani et al. 2010), Mitokeima (MBL life science), VDAC1 shRNA-puromycin, VDAC1-3XFLAG, VDAC1-GFP (1-10) and PHB2-GFP (11) (Vector builder). MitoBlue is a kind gift from Dr. Oishee Chakrabarti (Saha Institute of Nuclear Physics, Kolkata, India).

### *Drosophila* lines

Actin Gal4/Cyo is gifted by Dr. Pralay Majumder (Presidency University, Kolkata, India), UAS ATG8a mcherry is gifted by Prof. SC Lakhotia (Banaras Hindu University, Varanasi, India). UAS mitoKeima, Porin A2/Cyo and Porin RV are procured from Korea Drosophila Resource Center (KDRC) ^34^. These lines are maintained in standard cornmeal food. Rotenone / vehicle administration is done by mixing the proper concentration with the food (Kim et al. 2019).

### Cell culture and treatments

Human mid-brain derived dopaminergic neuroblastoma cell line SH-SY5Y or human embryonic kidney (HEK 293T) cells are cultured in DMEM (Gibco) with sodium bicarbonate (3.75 g/l), FBS (10%, Gibco), and penicillin-streptomycin (1%, Thermo) in a humidified incubator at 37 °C at 5% C02. Control and VDAC1 knockout HEK293T cells were procured from Abcam. Cell transfections are done using Transfectin TM (Biorad). Stable VDAC1 KD SH-SY5Y cell is obtained by transfecting VDAC1shRNA and selected against puromycin treatment following standard protocol. MitoBFP, VDAC1-GFP (1-10) and PHB2-GFP (11) were expressed in the mentioned cell lines for 36h. For measuring the raw IntDen we utilized freely available Image J software. For 3D rendering we used volume j plugin.

CCCP (10 µM) treatment, in presence or absence of MG-132 (50 µM) is given for different time points (1 to 8h), as mentioned in the relevant experimental procedure.

### Mitochondria isolation and treatments

Mitochondria were isolated from rat brain/cells/Drosophila by differential centrifugation following the protocol as mentioned previously ^45^. In brief, Cell/tissue is homogenized in mannitol–sucrose buffer (225 mM mannitol, 75 mM sucrose, 5 mM HEPES, 0.1 mM EGTA, pH 7.4, supplemented with 2% BSA) and then centrifuged at 1,500 × g (at 4°C for 6 min). The supernatant is again centrifuged at 7,000 × g (6 min). The pellet is washed with mannitol–sucrose buffer and re-suspended in appropriate amount of mannitol–sucrose buffer.

10 µM CCCP is treated for 20 min to induce depolarization, then crosslinked for another 30 min. Samples are immediately prepared in Laemmli buffer and processed for immunoblotting in mentioned conditions. The concentration of the crosslinkers are: bismaleimidohexane (BMH, Invitrogen)-5 mM, dithiobis-succinimidyl propionate (DSP)-2mM, ethylene glycol bis-succinimidyl succinate (EGS, Invitrogen) -50 µM for rat brain mitochondria and 1mM for isolated cell line mitochondria. In relevant experiments, 10 min prior treatment of 4,4′-diisothiocyanostilbene-2,2′-disulfonic acid (DIDS, 100 µM) is given.

### Protease (Trypsin) protection/digestion assay

For isolated mitochondria (from rat brain and Drosophila), around 50 µg is treated with either buffer or 200 μg/ml Trypsin (TPCK treated; Thermo) for 10 min at 4°C in Trypsin digestion buffer (10 mM sucrose, 0.1 mM EGTA/Tris and 10 mM Tris/HCl, pH 7.4); then, the samples are mixed with Laemmli buffer + β-mercaptoethanol and heated at 95°C.

For cells, we followed the protocol as described previously^45^. Briefly, cells from different genetic background are harvested after treatment (CCCP-10 µM, DIDS pretreatment-100 µM, 4h) and permeabilized (0.015% digitonin, 2 min), followed by washing with PBS. Cell pellet is re-suspended in trypsin digestion buffer and divided equally into two parts; out of which one received Trypsin (200 µg/ml). Both of the tubes are kept on ice for 5 min and then 2X Laemmli buffer + β-mercaptoethanol is added. Samples are separated in by SDS PAGE and processed for immunoblotting.

### Immunoblot and Immunoprecipitation

Crosslinked samples are resolved using SDS PAGE in non-reducing condition followed by Immunoblotting. For SDS-PAGE which is done under reducing condition, protein lysates are prepared in RIPA (supplemented with protease inhibitor cocktail, Thermo). Immunoblotting is carried out following the standard procedure. Secondary antibodies conjugated to HRP (Biobharati, India) were used (1:3,000) and visualized with ECL chemiluminescence (Invitrogen).

For second dimension SDS-PAGE, gel pieces are incubated with either β-marcaptoethanol or 0.5N NH_4_OH (as mentioned) for 1h, 37 °C in continuous shaking, followed by 20 min incubation with 2X Laemmli buffer. The gel pieces are kept on top of a resolving gel and sealed with agarose.

For third dimension SDS-PAGE, isolated gel pieces are further incubated with 0.5 NH_4_OH 1h, 37 °C and resolved again.

For immunoprecipitation, approximately 1 mg protein sample kept in lysis buffer (25 mM Tris HCL, pH 7.4, 150 mM NaCl, 1% NP40, 1 mM EDTA, 5% glycerol, 0.1% tween 20) and washed with equilibrated Protein A/G agarose beads (Santacruz biotechnology). Equal amount of cleared lysate is incubated with respective antibodies (1:100), 4 °C overnight, and on the next day Protein A/G agarose beads are co-incubated for 2h at room temperature. Afterwards beads are washed repeatedly and protein samples are eluted using 2X Laemml buffer(+ β mercaptoethanol).

### Evaluation of mitokeima in cells and Drosophila wing

Cells are transfected with mitokeima and after 48h, CCCP treatment is given to induce mitophagy (8h). A separate group also received DIDS pretreatment (100 µM). Signal intensity from single cell suspension is determined by FACS. High mitophagic cells ^46^ are quantified by Flowjo software.

For Drosophila, mitokeima expressing live fly wings are visualized under confocal microscope (Zeiss, LSM980). For all the experiments Ex: 440 / 568 and Em: 610 is used. For thorax: thoracic muscles are dissected out on coverslip and kept in PBS (pH 7.4) while imaging.

### Immunofluorescence and confocal imaging

Fixed cells (4% PFA) on coverslips are permeabilized with 0.1% triton X-100 and blocked with 2% BSA. For Drosophila thoracic muscle, samples are fixed in 4% PFA for 3days and transferred to 30% sucrose. Cryosections on slides are permeabilized with 0.2% Triton X-100. BSA (8%, 1h) is used for blocking. Samples are incubated with specific primary antibodies at 4◦C overnight in a humid chamber and tagged with Alexa fluor conjugated secondary antibody. Samples are mounted with anti-fade reagent (Invitrogen).

For Duo link-proximity ligation assay, 4%PFA fixed samples are blocked and incubated with primary antibodies (rabbit-PHB2 and mouse-LC3), as described above. A kit is used to develop the signal following manufacturers protocol (Sigma Aldrich).We visualized the signal by using a high resolution confocal microscope (Zeiss LSM980).

### Transmission electron microscopy

To acquire TEM images, we followed the protocol as stated previously (Chakraborty et al. 2018).

### Statistics

Bar graphs represented in the figures are mean ± SEM. Two-tailed Student’s t-test (or mentioned otherwise) or one-way ANOVA is used to decide the level of significance. Relevant post hoc tests are mentioned along with the figure legend. P ≤ 0.05 was considered as significant difference.

## Acknowledgement

We are thankful to Science and Engineering Research Board (SERB) for financially supporting the study. We also thank International Brain Research Organization (IBRO) for supporting international travel expenses under “Young IBRO Regions Connecting Award”. CIF division is acknowledged for instrumentation support. MR and RM is recipient of senior research fellowship from the Council of Scientific and Industrial Research - (CSIR), India. CB is a recipient of a senior research fellowship from the University Grants Commission (UGC), India. SN is financially supported by SERB funded project GAP410.

## Author contribution

JC designed the experiments, analyzed data and wrote the manuscript. MR and SN performed most of the experiments. CB contributed for mitokeima related quantification. CB and RM performed some of the cell and *Drosophila* related immunoblots. EM, EZ and FC performed TEM experiments. EZ critically evaluated the manuscript. All authors discussed and contributed to the manuscript.

## Competing interest

The authors declare no competing interests.

## List of abbreviations

Prohibitin 2: PHB2
Voltage-dependent anion-selective channel protein 1: VDAC1
PTEN induced kinase1: PINK1
outer mitochondrial membrane: OMM
inner mitochondrial membrane: IMM
Gene Expression Omnibus: GEO
Substantia nigra: SN
kilodaltons: kDa
sodium dodecyl sulfate–polyacrylamide gel electrophoresis: SDS-PAGE
Carbonyl cyanide m-chlorophenyl hydrazone: CCCP
Transmission electron microscope: TEM
Knockout: KO
Knockdown: KD
dithiobis-succinimidyl propionate: DSP
ethylene glycol bis-succinimidyl succinate: EGS
bismaleimidohexane: BMH
Human embryonic kidney 293 cells: HEK393T
,4′-diisothiocyanostilbene-2,2′-disulfonic acid: DIDS
Porin KO: A2/A2
Porin reintroduced: RV
wild type: WT
systemic lupus erythematosus: SLE

## Notes

### Competing Interest Statement

The authors have declared no competing interest.

