## Supplementary figures and images for "Efficient Prohibitin 2 exposure during mitophagy depends on Voltage-dependent anion-selective channel protein 1"

### Figure S1

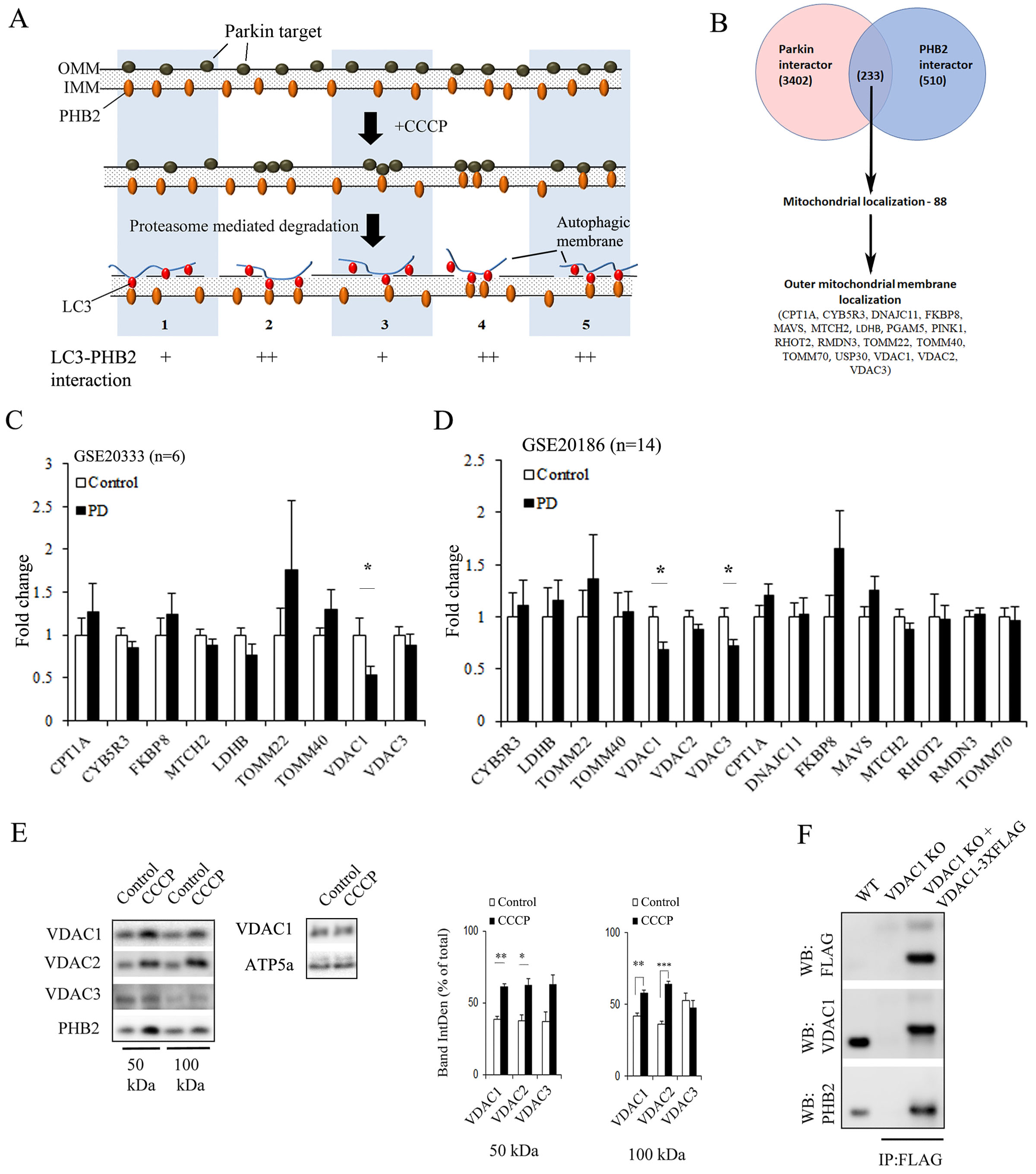

### Figure S2

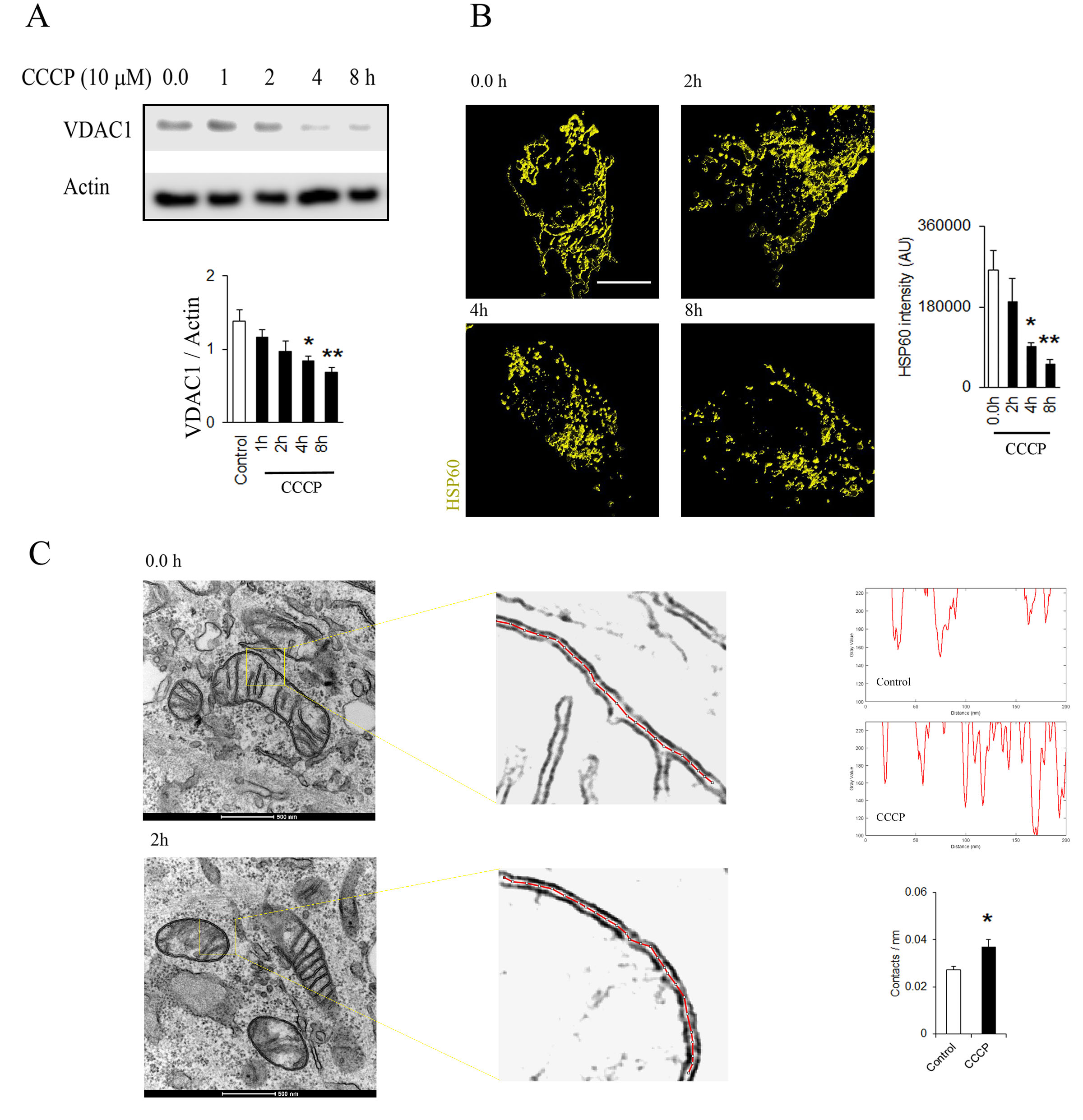

### Figure S3

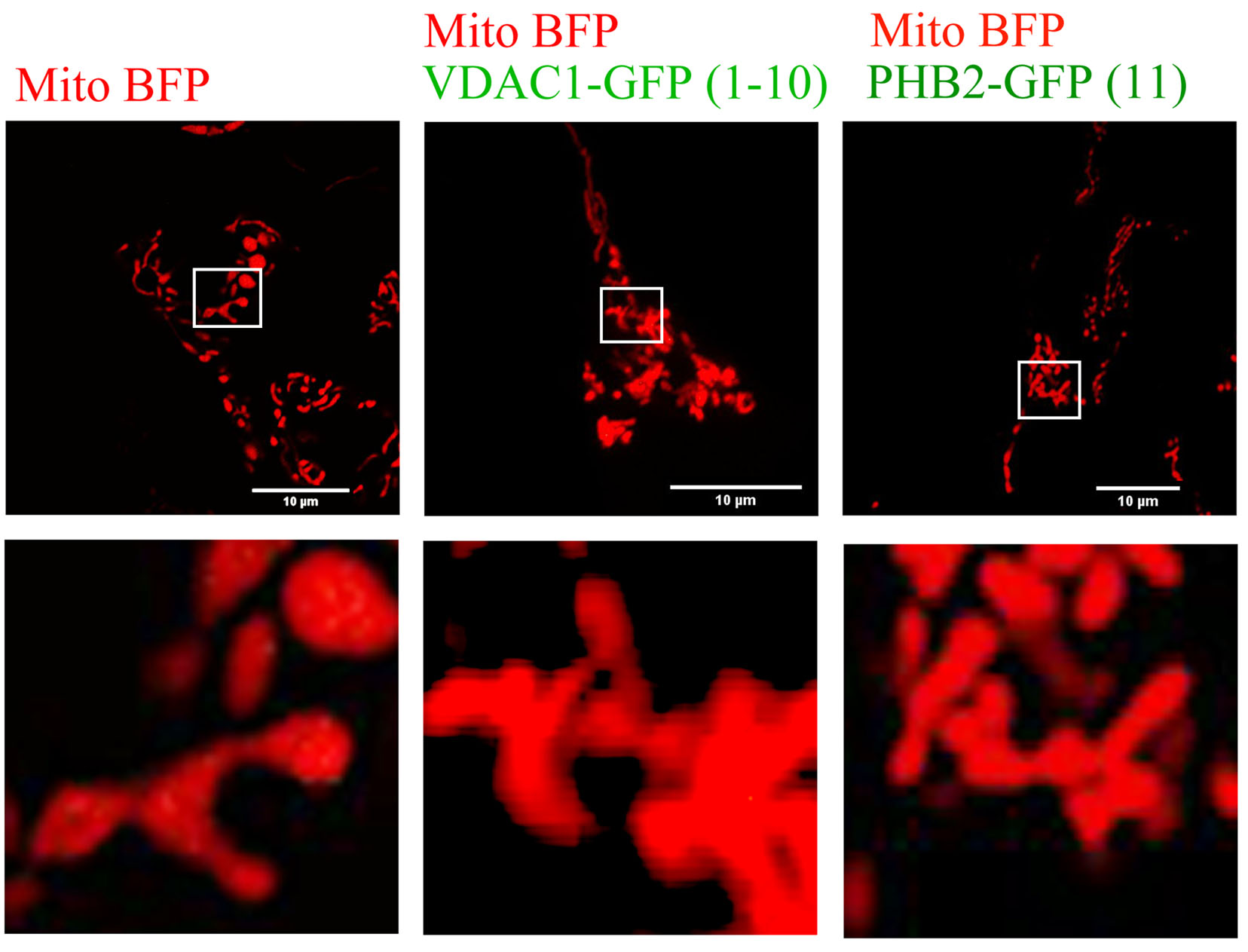

### Figure S4

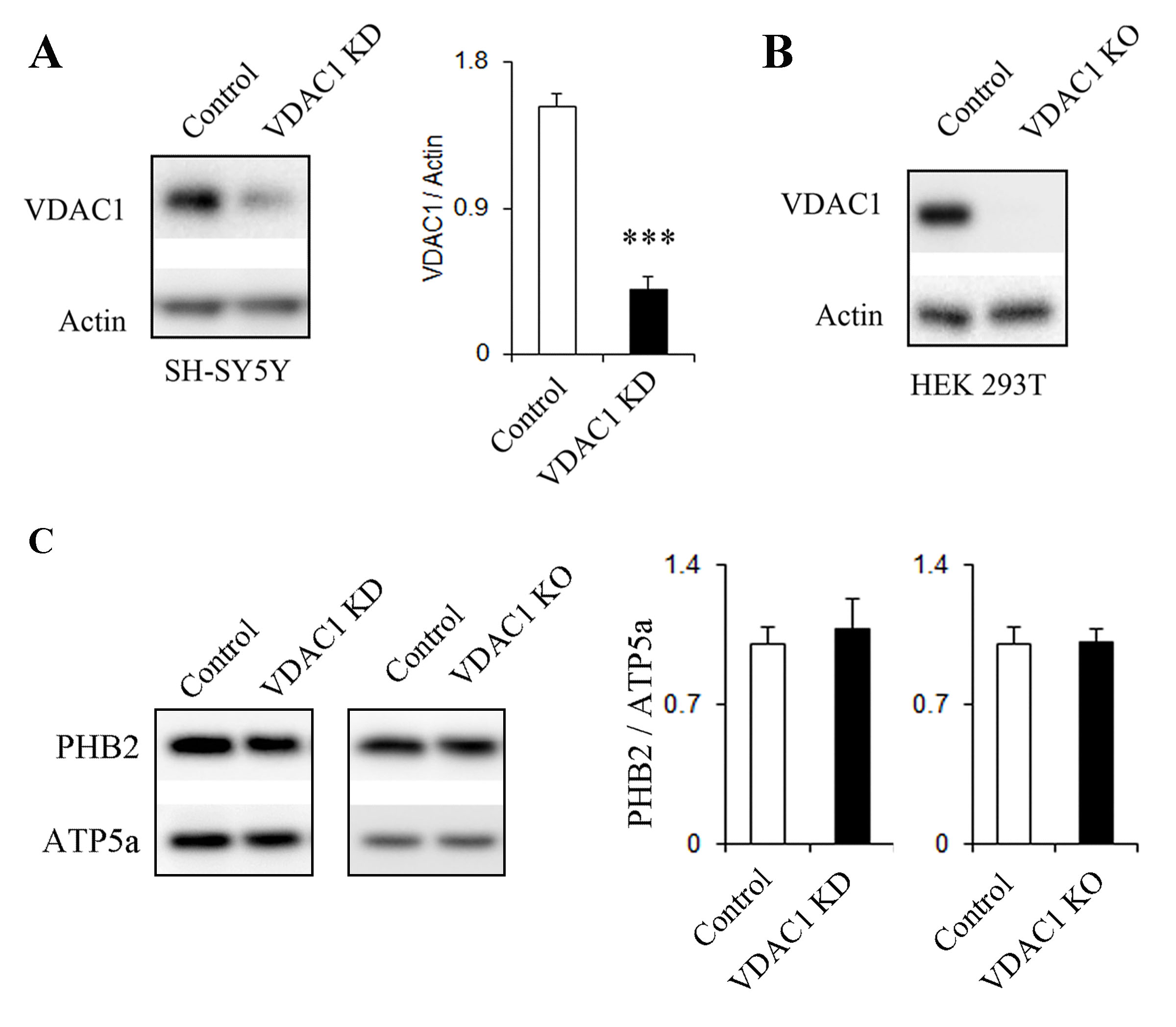

### Figure S5

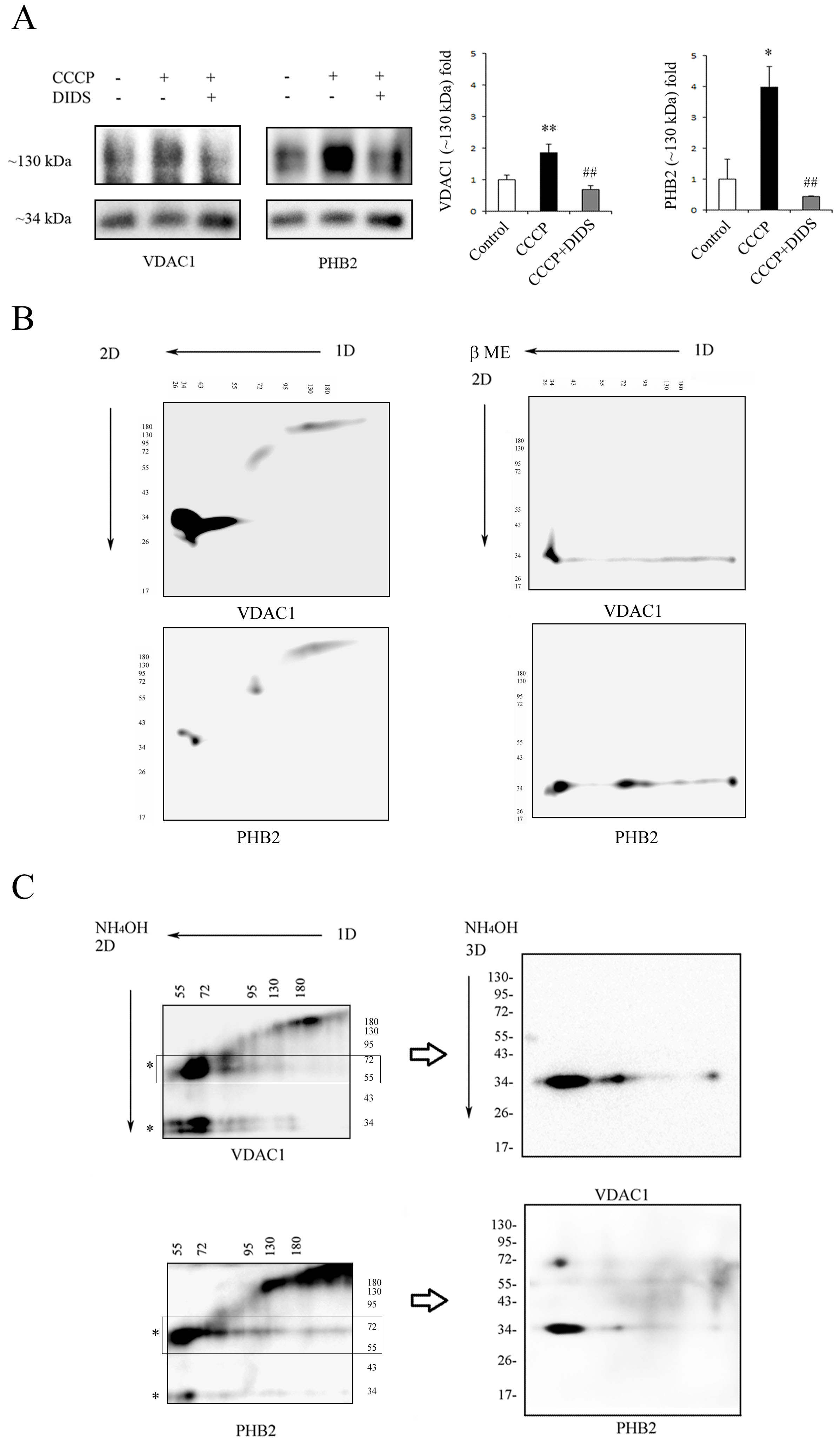

### Figure S6

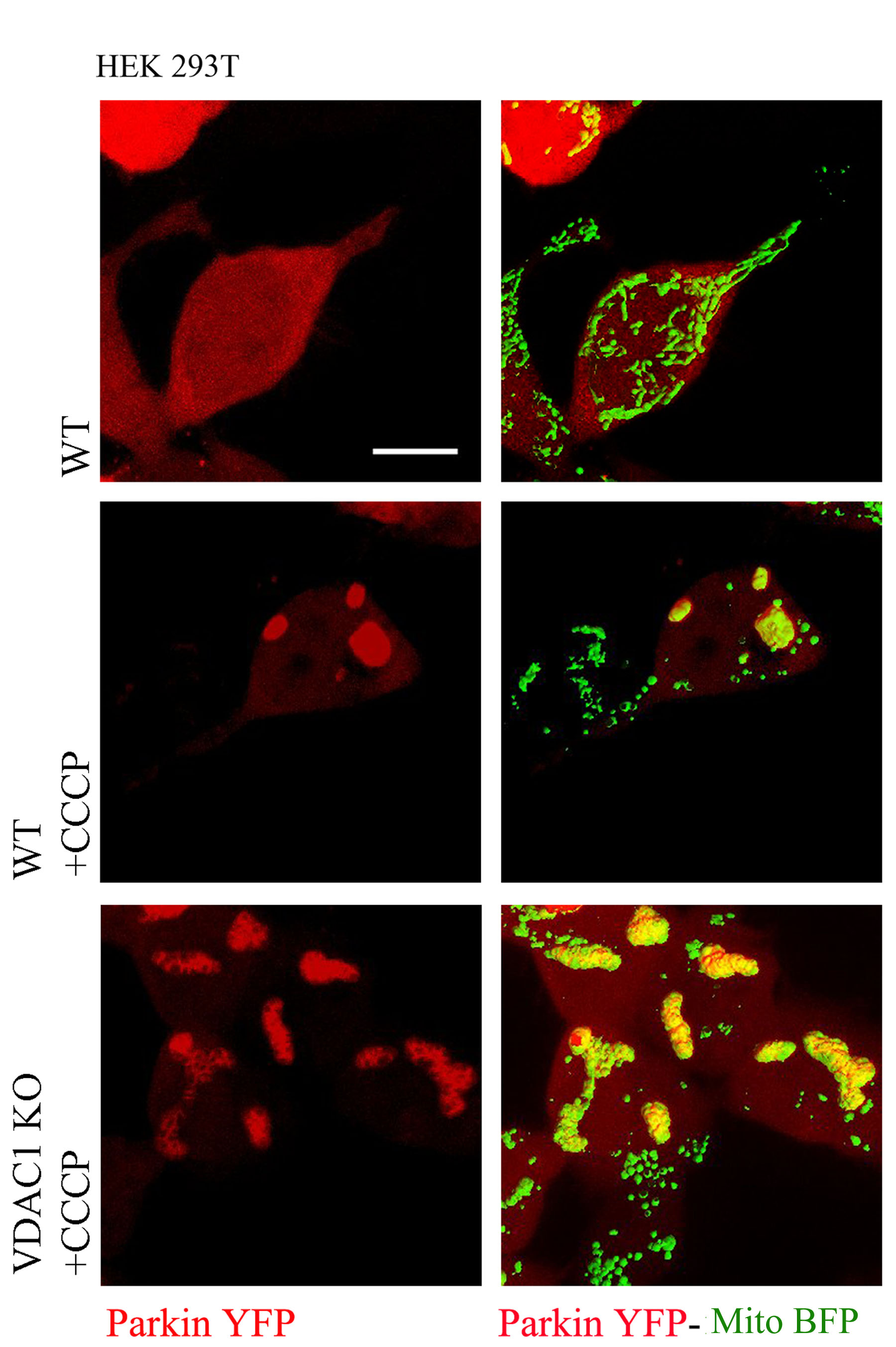

### Figure S7

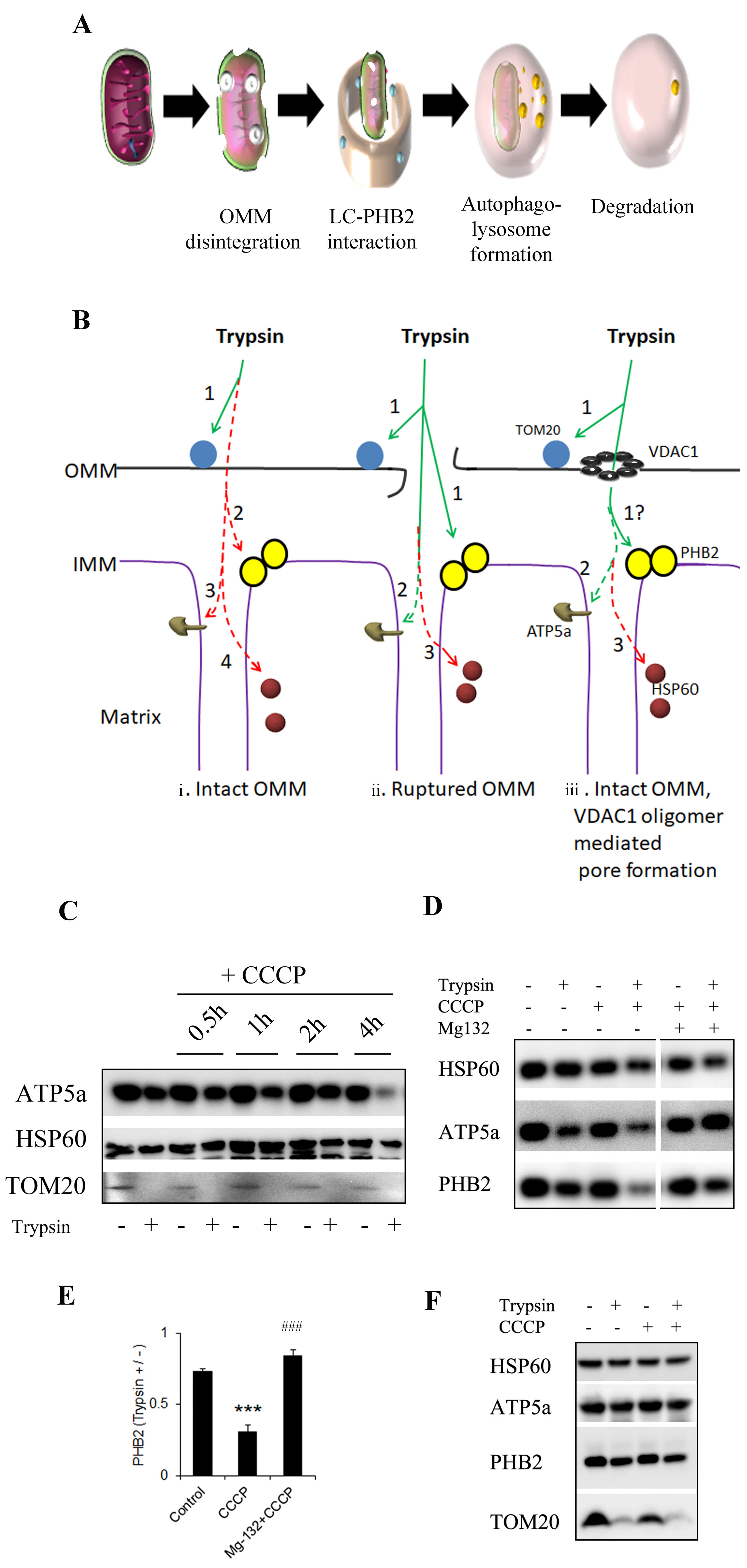

### Figure S8

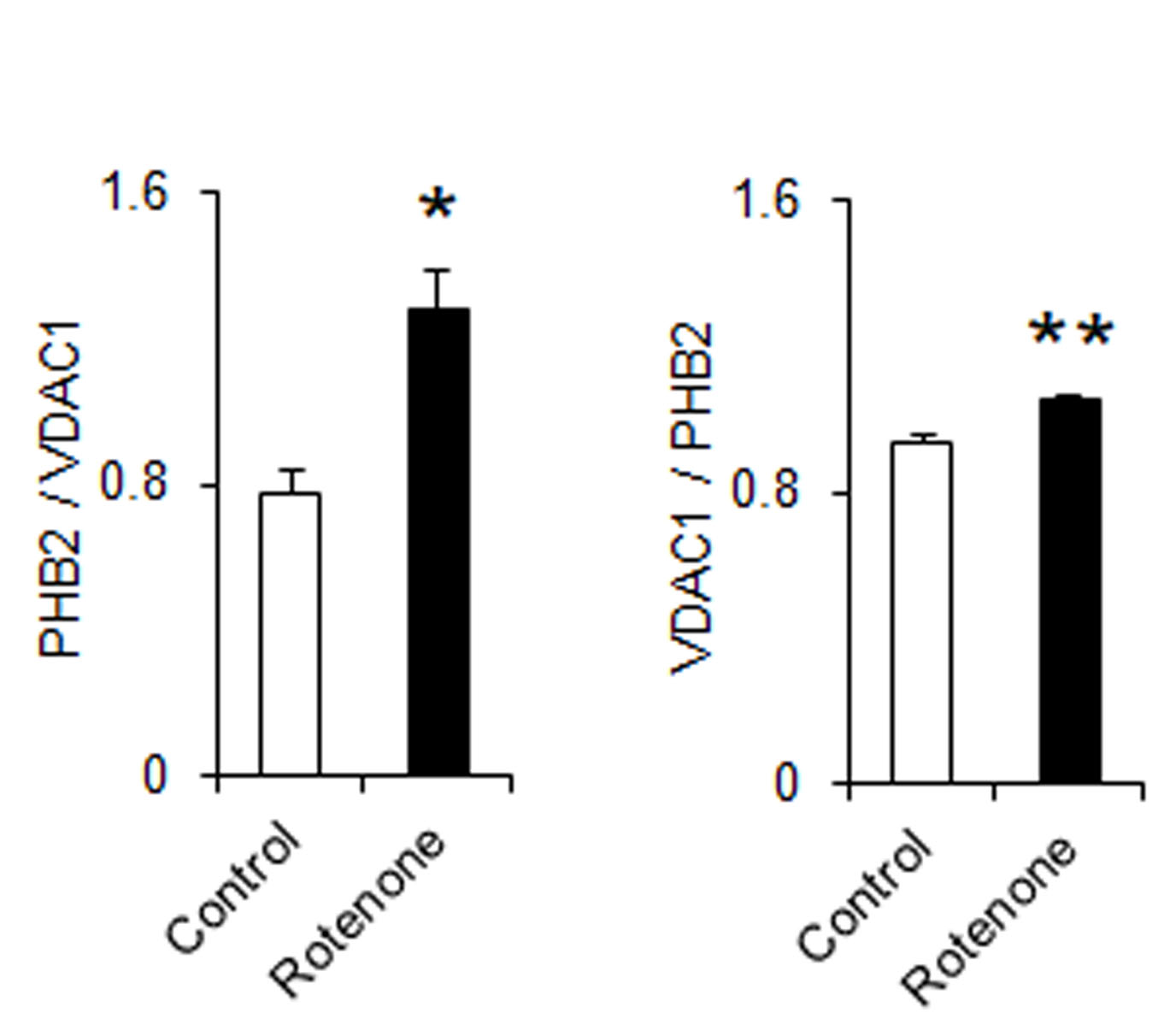

### Figure S9

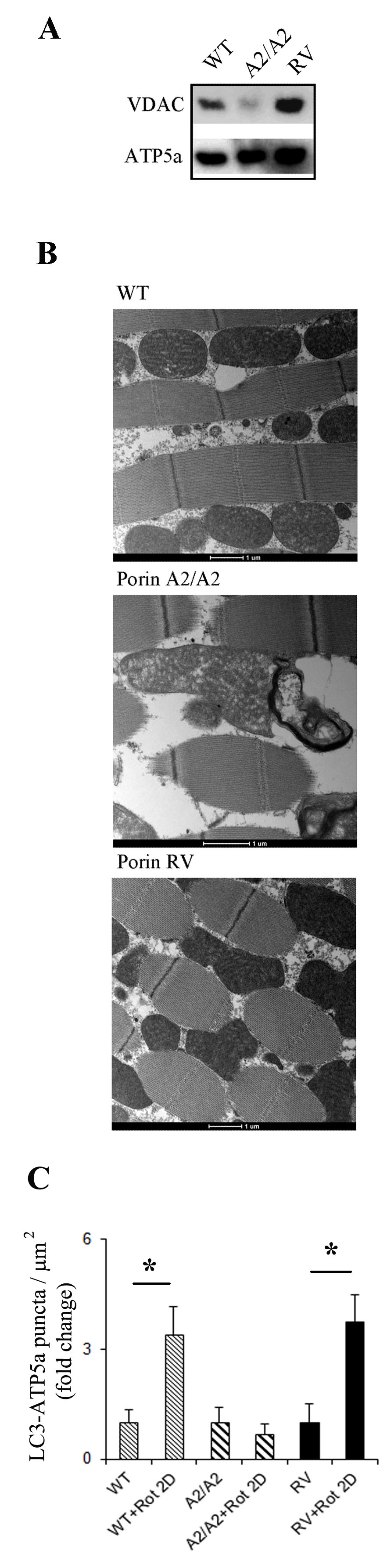
