## Supplementary figure legends for "Efficient Prohibitin 2 exposure during mitophagy depends on Voltage-dependent anion-selective channel protein 1"

**Supplementary Figure legend**

**Figure S1. Hypothesis and identification of VDAC1.**

**A.** Graphical image demonstrates the hypothetical distribution of outer mitochondrial (OMM) and inner mitochondrial membrane (IMM) proteins before and after depolarization in five different combinations (mentioned as 1-5 in columns). Black or orange circles represent Parkin substrate or PHB2 respectively. Proteasome mediated degradation of OMM proteins may lead to different level of interaction (mentioned as “+” or “++”) between LC3 (mentioned as red circles on autophagic isolation membrane-blue line) and PHB2.

**B.** Image demonstrates that there are 233 common interacting partners of Parkin and PHB2, out of which 88 reside on mitochondria. Out of these, only 18 are OMM proteins and thus may be useful to verify the above depicted scenario.

**C.** and **D**. Expression values for control / Parkinson’s disease substantia nigra brain region were obtained from NCBI transcriptome database and analyzed by Gene Expression Omnibus (GEO). Bar graphs represent mean fold change ± SEM. *P ≤ 0.05. GEO accession number : C - GSE20333 (n=6); D - GSE20186 (n=14). Student’s t test.

**E.** Isolated rat brain mitochondria are treated with CCCP (10 µM, 20 min) or DMSO (control) and protein lysates are separated through 100 or 50 kDa cut off protein concentrators. Equal volumes of protein are subjected for immunoblotting. A portion of isolated mitochondria (± CCCP) were directly processed for immunobloting to determine the levels of VDAC1 and ATP5a in the different treatment groups. Bar graphs represent mean ± SEM. *P ≤ 0.05, **P ≤ 0.01, ***P ≤ 0.005, n=3. Student’s t test.

**F.** VDAC1 KO HEK cells are transfected with empty or VDAC1-3XFLAG plasmid and isolated mitochondrial lysate is subjected to immunoprecipitation for FLAG (IP: FLAG). Protein samples from WT HEK cells were separated simultaneously to detect the shift of VDAC1 signal.

**Figure S2. Time dependent reduction of VDAC1 level and mitochondrial mass after CCCP treatment.**

**A.** Representative immunoblot of VDAC1 from SH-SY5Y cells. Time points indicate CCCP treatment for different periods. Actin is considered as a loading control. Bar graphs represent mean ± SEM. **P ≤ 0.01, n= 3, One way ANOVA followed by Dunnett’s test.

**B.** Mitochondrial elimination in SH-SY5Y cells after CCCP treatment is evaluated by HSP60 immuno-staining. Scale bar : 10 µm. Bar graphs represent mean intensity ± SEM. *P ≤ 0.05, **P ≤ 0.01, at least 30 cells were evaluated. N=3. One way ANOVA followed by Dunnett’s test.

**Figure S3. Expression of VDAC1-GFP (1-10) and PHB2-GFP (11) in mitoBFP expressing HEK 293T cells.**

VDAC1 KO HEK293T cells were transfected with either mitoBFP+ VDAC1-GFP (1-11) or mitoBFP+PHB2-GFP(11). Representative images show undetectable GFP signal. Scale bar 10 µM.

**Figure S4. Characterization of stable VDAC1 knock down or knock out cells**

**A-B.** Validation of stable VDAC1 knockdown or knockout in SH-SY5Y and HEK293T cells, respectively. Actin is used as loading control. Bar graphs represent mean intensity ± SEM. ***P ≤ 0.005. Student’s t test, n=3.

**C.** Mitochondrial PHB2 level in VDAC1 KD (SH-SY5Y) / KO (HEK) cells. Bar graphs represent mean intensity ± SEM, n=3.

**Figure S5. Effect of DIDS treatment on VDAC1-PHB2 complex formation during mitochondria depolarization and immunoblots for VDAC1 and PHB2 after second /third dimension SDS PAGE of DSP/EGS crosslinked mitochondrial samples.**

**A.** Isolate rat brain mitochondria are treated as depicted in Figure 1A, except another group is incorporated where DIDS (100 µM) is co-incubated with CCCP.

Bar graphs represent mean fold change (± SEM , n=3). *P ≤ 0.05, **≤ 0.01 (compared to control), # P ≤ 0.05 (compared to CCCP treated group). One way ANOVA followed by Tukey’s multiple comparison test.

**B.** Isolated rat brain mitochondria are treated with CCCP (10 µM, 20 min) and crosslinked with DSP. After first dimention (1D) SDS-PAGE, gel pieces are treated with 4X Laemmli buffer (+/- β - marcaptoethanol) for 30 min at 37˚C. Subsequently a second dimension (2D) SDS PAGE is performed. Arrows indicate the direction of the run.

**C.** 2D analysis of EGS crosslinked samples, as depicted in B, except gel pieces were reduced by NH_4_OH. Pieces were cut from ~70 kDa (marked by boxes) and a 3D analysis were performed after further reduction.

**Figure S6. Parkin localization in HEK cells with / without CCCP treatment.**

Parkin-YFP and mito-Blue protein expressing WT and VDAC1 KO HEK cells are treated with CCCP (8h). Images are taken to determine Parkin localization after depolarization. Images are representative of 3 different experiments. Scale bar 10 µm.

**Figure S7. Evaluated events during mitophagy and mitochondrial outer membrane integrity after CCCP treatment.**

**A.** Events that are evaluated to assess mitophagy level in the study are mentioned in the graphical image.

**B.** Differential sensitivity of HSP60, ATP5a, PHB2 and TOM20 in three different scenarios which were assessed for the study is depicted. Green arrows indicate readily accessible by Trypsin, whereas red dotted arrows specify proteins which are less likely to be degraded by the same.

**C.** SH-SY5Y cells are treated with CCCP (10 µM) for different time periods and after permeabilization, proteolysis of ATP5a and HSP60 by Trypsin are evaluated by immunoblotting.
**D-E.** SH-SY5Y cells are treated with CCCP for 4h in presence / absence of MG-132 (50 µM). Sensitivity of the represented proteins towards Trypsin was evaluated by immunoblotting. Bar graphs (E) represent mean intensity ± SEM. ***P ≤ 0.005, ### P ≤ 0.005 compared to CCCP . Student’s t test, n=3.

**F.** Isolated rat brain mitochondria are treated with CCCP for 20 minutes and sensitivity of PHB2, ATP5a, HSP60 and TOM20 towards Trypsin digestion is evaluated by immunoblotting. Blots are representative of three different experiments.

**Figure S8. Quantification for Figure 3E.**

Bar graphs represent mean ± SEM. **P ≤ 0.05, **P ≤ 0.01when compared to rotenone treated group . Student’s t test, n=3.

**Figure S9. VDAC level, mitochondrial ultrastructure and quantification LC3-ATP5a localization in the fly lines used.**

**A. Porin (**VDAC) levels in isolated mitochondria from the indicated *Drosophila* lines.

**B.** Transmission electron microscopic images of fly thoracic muscle mitochondria are represented for comparative assessment. Scale bar is mentioned in the images.

**C.** Quantification of LC3-ATP5a puncta overlap of Figure 4C. Student’s t test. *P ≤ 0.05 when compared to the respective non treated group. n= 4-6.
